# Tuning into xanthan: A conserved yet flexible polysaccharide utilization system in *Microbacterium*

**DOI:** 10.64898/2026.04.20.719765

**Authors:** Bianca Laker, Michael Thomas, Wiebke Weber, Prisca Viehoever, Anja Meierhenrich, Levin J. Klages, Tobias Busche, Karsten Niehaus, Andrea Bräutigam, Marion Eisenhut

**Affiliations:** Computational Biology, Faculty of Biology, Bielefeld University, Bielefeld, Germany; Medical School OWL, Bielefeld University, Bielefeld, Germany; Center for Biotechnology (CeBiTec), Bielefeld University, Bielefeld, Germany; Proteome and Metabolome Research, Faculty of Biology, Bielefeld University, Bielefeld, Germany; Genetics and Genomics of Plants, Faculty of Biology, Bielefeld University, Bielefeld, Germany

**Keywords:** xanthan, Microbacterium, polysaccharide utilization, regulation, microbial ecosystems, evolution

## Abstract

Bacteria encounter structurally complex extracellular polysaccharides in natural environments, yet the regulatory and evolutionary basis of their utilization remains poorly understood. Here, we isolated a soil-derived *Microbacterium* strain, named *Microbacterium xanthanicum* UB-LE1, that grows on xanthan as the sole carbon source. We dissected the genetic and regulatory architecture underlying this capability. Genome sequencing combined with transcriptomic and proteomic profiling uncovered a discrete, strongly inducible regulon associated with xanthan utilization, encoding 23 proteins with five secreted proteins and three candidate transcriptional regulators. DNA-affinity purification sequencing confirmed two regulators binding to operons within the *xanthan utilization* locus. Comparative genomics across the Microbacteriaceae revealed conserved and lineage-specific features of this system and supports recent acquisition and modular integration of the locus, with at least two predominant architectural variants possibly shaped by substrate availability and ecological specialization. Coordinated induction at both the transcript and protein levels, together with two experimentally validated regulators, points to tight regulatory control of complex polysaccharide degradation in *Microbacterium xanthanicum* UB-LE1.

Together, these findings provide mechanistic and evolutionary insight into how bacteria adapt to complex extracellular carbohydrates, expand current knowledge of xanthan turnover in microbial ecosystems, and establish a framework for exploring the emergence and diversification of specialized polysaccharide utilization pathways across bacterial taxa.

**IMPORTANCE:** Microorganisms are central drivers of carbon turnover in soils and other terrestrial ecosystems, determining the availability of nutrients and shaping microbial community structure. A significant portion of soil carbon is contained in extracellular polysaccharides, yet the pathways by which microorganisms degrade these complex polymers remain poorly understood. Xanthan, a structurally complex and widely produced microbial exopolysaccharide, represents a persistent and largely overlooked carbon pool. By dissecting the genetic, regulatory, and evolutionary basis of xanthan utilization in *Microbacterium xanthanicum* UB-LE1, this study advances our understanding of how soil bacteria adapt to complex extracellular carbohydrates and how substrate availability shapes the emergence and diversification of specialized metabolic pathways. Importantly, the identification of additional xanthan-active enzymes and regulatory components in *M. xanthanicum* UB-LE1 opens opportunities for targeted modification of xanthan structure and properties, paving the way for new biotechnological applications in food, materials, and industrial biotechnology, while linking microbial ecology to functional innovation.

## INTRODUCTION

Microorganisms play a central role in shaping carbon turnover in soils and other terrestrial ecosystems through the secretion and degradation of extracellular polymers. Among these polymers, exopolysaccharides (EPS) are secreted into the environment to form biofilms, which provide structural support for microbial communities and act as carbon sources for specialist degraders. Understanding how microorganisms sense, transport, and metabolize structurally complex EPS is essential to uncovering both ecological interactions and the evolution of specialized metabolic pathways.

Xanthan is a high-molecular-weight, anionic heteropolysaccharide produced by members of the *Xanthomonas* genus. It consists of a β-(1,4)-glucan backbone substituted with trisaccharide side chains containing acetylated or pyruvylated mannose and glucuronic acid. In plant-pathogenic *Xanthomonas campestris*, xanthan contributes to virulence by suppressing plant immune responses, protecting the bacteria from host-derived and environmental stresses, and facilitating adhesion, aggregation, and colonization of the apoplast (1–3). Loss of xanthan biosynthesis attenuates virulence, highlighting its central role in host–pathogen interactions. Beyond its role in plant pathogenesis, xanthan persists in soil for extended periods, with portions reported to resist degradation for several months (4) due to its high molecular weight and complex chemical structure, providing a stable carbon pool for microbial communities.

The structural complexity of xanthan has also driven its widespread industrial use. Its high viscosity and shear-thinning properties make it a ubiquitous additive in food, cosmetics, personal care, and industrial formulations (5, 6). Emerging applications in biomedical engineering (7) and geoengineering (4) highlight the growing relevance of understanding xanthan metabolism and biodegradation. Because it is biodegradable, identifying microbial enzymes that can modify or degrade xanthan has both ecological and biotechnological significance, offering potential avenues for tailored modification of xanthan properties for novel applications. Despite the industrial and ecological relevance of xanthan, the mechanisms by which fungi and bacteria degrade this polysaccharide (8) are only partially understood. Xanthan degradation typically begins with an extracellular xanthan lyase (PL8) that liberates terminal pyruvylated mannose residues, followed by an endoxanthanase (GH9 or GH5 family) that cleaves the backbone into tetrasaccharides (8). Complete turnover requires intracellular enzymes to remove acetyl groups and cleave the resulting oligosaccharides into monosaccharides (8, 9), as well as transport systems to import these products and regulatory proteins to coordinate expression of the catabolic machinery. A variety of xanthan-degrading bacteria have been isolated from diverse environments including soil (10), sludge (11), and mammalian intestines (8). These include *Bacillus* GL1 (12) and *Microbacterium* XT11 (13), both of which possess secreted PL8 xanthan lyases, membrane-bound GH9 enzymes, and intracellular GH38 and GH3 enzymes (10), and also include *Paenibacillus nannanis* (9, 14, 15), *Bacteroides* and *Ruminococcus* (8), *Cohnella* sp. 56 (VKM B-36720) (16), and *Paenarthrobacter ilicis* (17). While many xanthan-degrading enzymes are known, information on their regulatory control is scarce. In other extracellular polysaccharide systems, catabolic genes are frequently organized into co-regulated clusters. For instance, *polysaccharide utilization* (*pul*) loci in human gut Bacteroidetes comprise genes for extracellular enzymes, transporters, intracellular enzymes, and regulators (18). Secretion of large or high-molecular-weight proteins is energetically costly, and these systems are typically under tight transcriptional control (18). Similarly, gene clusters for pectin degradation in *Erwinia chrysanthemi* (19) or rhamnogalacturonan degradation in *Bacillus subtilis* (20) are regulated by specific transcription factors or two-component systems. However, no such regulatory frameworks have been comprehensively described for xanthan utilization.

To investigate how bacteria adapt to structurally complex extracellular carbohydrates, we focused on a newly isolated, soil-derived *Microbacterium* strain capable of using xanthan as its sole carbon source. This strain, *M. xanthanicum* UB-LE1, provides a unique system to connect ecological persistence of xanthan with the microbial strategies required for its degradation. By integrating genomic, transcriptomic, and proteomic data, we map the functional components involved in xanthan utilization, including secreted enzymes, intracellular catabolic proteins, and regulatory elements coordinating their expression. Comparative analyses across related Microbacteria indicate that these pathways combine both conserved mechanisms and lineage-specific adaptations, reflecting evolutionary pressures imposed by substrate availability and ecological specialization. Studying *M. xanthanicum* UB-LE1 thus illuminates how bacteria detect, import, and metabolize complex polysaccharides in natural environments, while also providing a framework for exploiting these systems in biotechnology.

## RESULTS

### Isolation of *Microbacterium xanthanicum* UB-LE1 as xanthan-degrading bacterium

To isolate a naturally occurring xanthan-degrading bacterium, a topsoil sample from the eastern Teutoburger Forest in Bielefeld, Germany, at approximately 52,01387° N, 8,48334° E was enriched for capable xanthan degraders. After several rounds of single colony restreaking, a single orange colony was isolated and provisionally named UB-LE1. Nanopore sequenced DNA assembled into a full genome as a single contig with a total size of 3,410,522 bp, a GC content of 70.46%, and BUSCO completeness of 99.2% (details of genome assembly and annotation in Laker et al. (Companion paper submitted to MRA). Since UB-LE1 showed the highest average nucleotide identity (ANI) values of 98.22% with *Microbacterium* sp. Leaf436 (Table S2) and TYGS classified UB-LE1 as closest to *Microbacterium enclense* NIO-1002 (details of phylogenetic analysis in Laker et al. (Companion paper submitted to MRA)), the strain was designated *Microbacterium xanthanicum* UB-LE1 (in the following *M. xanthanicum* UB-LE1).

*M. xanthanicum* UB-LE1 is a Gram-positive, rod shaped (1 µm by 3 to 4 µm) bacterium that lacks a flagellum and is non-motile (Fig. S1A). Its growth optimum was detected at 30 °C, pH 7, without addition of NaCl (Fig. S1B-D). Growth experiments in replicates identified *M. xanthanicum* UB-LE1 as an efficient user of xanthan as the sole carbon source with growth rates on xanthan of µ = 0.0653 + 0.0185 h^-1^ in comparison to glucose with µ = 0.102 + 0.004 h^-1^ (Fig. 1).

**Fig. 1.**
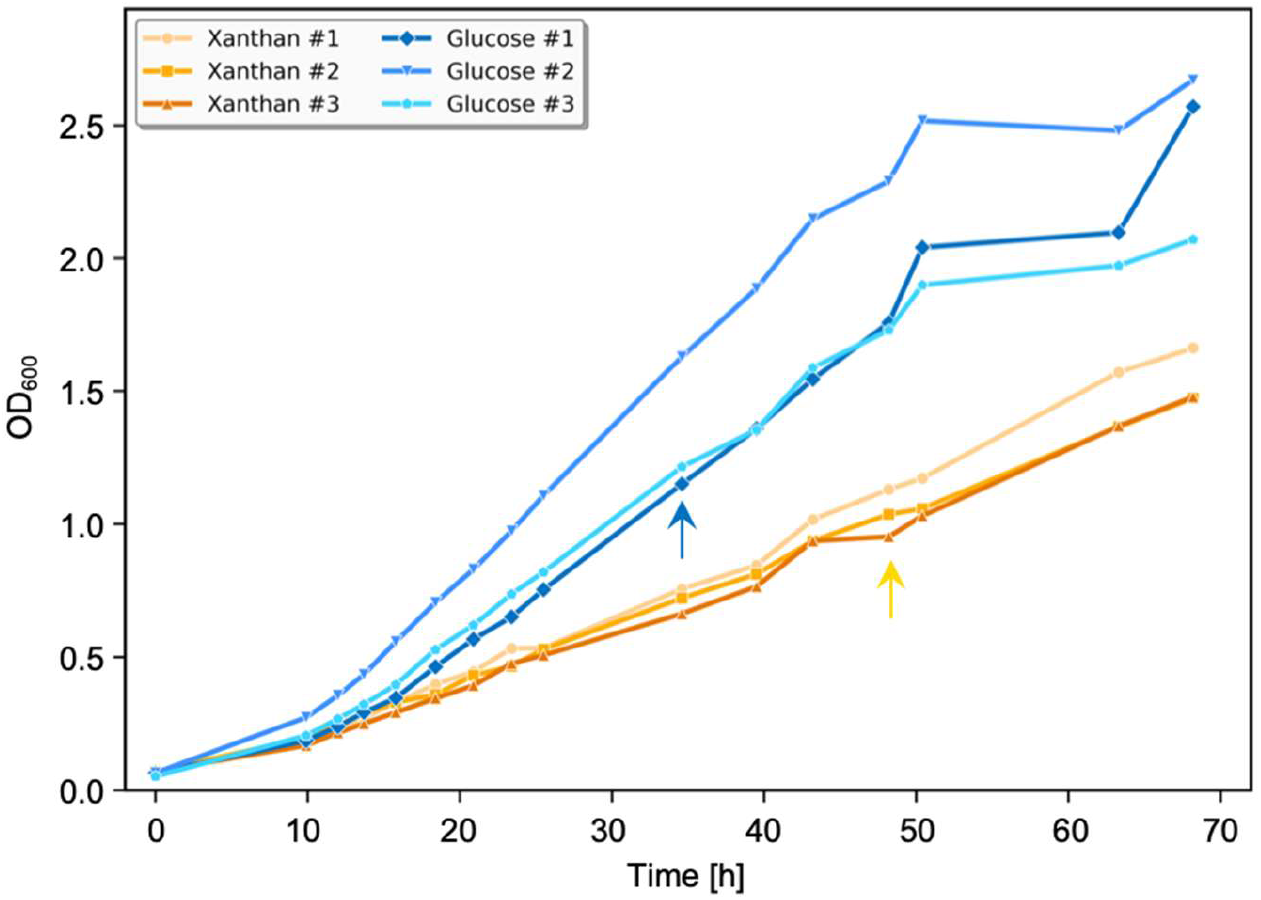
Growth and sampling of *M. xanthanicum* UB-LE1 for OMICS studies. Growth of M. *xanthanicum* UB-LE1 in M9 medium with either glucose or xanthan as C-source was performed on a shaker at 30 °C and 180 rpm. Sampling time points for transcriptomic and proteomic analyses are indicated with arrows (blue: glucose sampling; yellow: xanthan sampling). Three biological replicates were grown under each condition and sampled.

### Response to xanthan on transcript and protein level

To gain a comprehensive view on *M. xanthanicum’s* response to xanthan we performed a comparative OMICS approach. Samples for transcriptomic and proteomic (cellular and extracellular samples) analyses were taken after 34 h (+ glucose) and 48 h (+ xanthan), respectively, in the exponential grow phase and at a similar cell density (Fig. 1).

The transcriptome analysis detected 3,085 transcripts with xanthan as the carbon source and 3,093 transcripts with glucose as the carbon source (Table S1: mothertable). Samples were well separated in a principal component analysis (PCA) with 49% variation represented in the first principal component indicating the major source of variation in the transcriptome experiment is indeed the carbon source (Fig. 2A). The proteome analysis of the cellular protein detected 1,285 proteins with xanthan as the carbon source and 1,274 proteins with glucose as the carbon source (Table S1: mothertable). Samples were well separated in a PCA analysis with 51% of variation represented in the first component (Fig. 2B). The genome contained 202 genes with products that were predicted to be secreted by the secretion protein (SP) or twin-arginine translocation (TAT) pathways (Table S1: mothertable). Seventy-nine of these predicted extracellular proteins were detected in the supernatant with xanthan as the carbon source and 56 proteins were detected with glucose as the carbon source. The extracellular proteome samples carried higher variation as only 38% of variation in the first principal component separated both groups (Fig. 2C).

**Fig 2.**
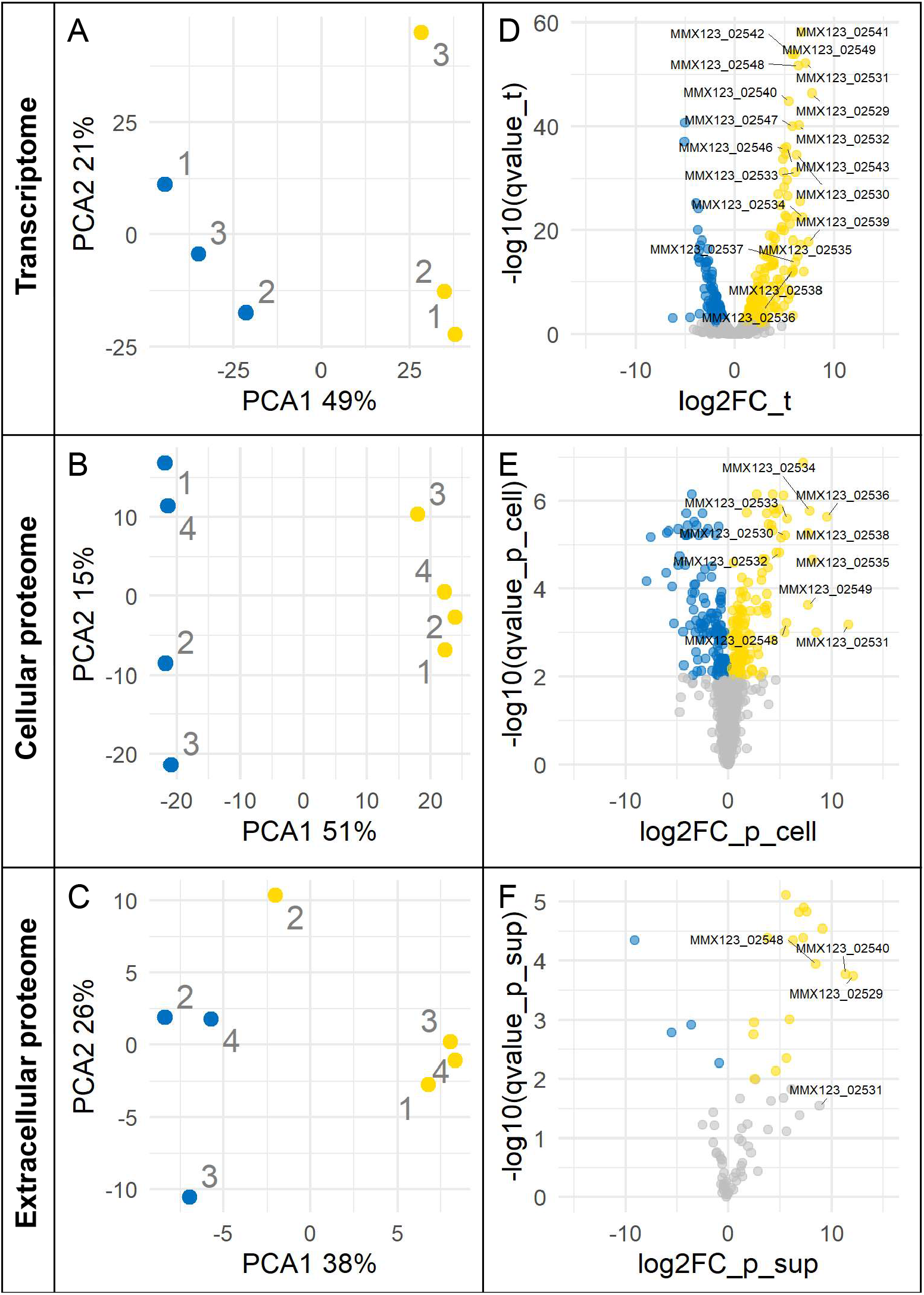
RNA-seq and proteomics analyses. Profiles of *M. xanthanicum* UB-LE1 grown under two different carbon source conditions. Principal component analysis of (A) transcriptome, (B) cellular proteome, and (C) extracellular proteome profiles from *M. xanthanicum* UB-LE1 grown on glucose or xanthan as the sole carbon source. Yellow points represent samples grown on glucose medium and blue points represent samples grown on xanthan medium. Biological replicates are marked by their labels. Volcano plot showing differential gene expression (D), differential cellular proteome (E) accumulation and differential extracellular proteome (F) accumulation between xanthan- and glucose-grown cells. Each point represents a gene or protein. Not significantly regulated events are colored in gray, significantly xanthan up-regulated events are colored in yellow, significantly glucose up-regulated events are colored in blue.

Statistical analysis detected 257 transcripts with significantly (*q* < 0.01) higher abundance in xanthan-dependent growth compared to glucose and 186 transcripts with significantly lower abundance in xanthan, that is higher abundance in glucose-dependent growth (Fig. 2D). When the 34 transcripts with at least 32-fold induction in xanthan were examined, they included *MMX123_00115, MMX123_00472* and *-473, MMX123_01228, MMX123_01822* and *-1823*, genes in a genome region from gene *MMX123_02529* to gene *MMX123_02549*, genes in a genome region from gene *MMX123_02601* to *MMX123_02608*, and genes in a genome region from *MMX123_03133* to *MMX123_03136* (Fig. 2D). Among the proteins, statistical analysis detected 149 cellular proteins as higher abundant in xanthan compared to glucose and 144 cellular proteins as higher abundant in glucose (Fig. 2E, Table S1: mothertable). When the proteins with fold-changes greater than 32-fold were examined, they included the protein encoded by *MMX123_00473*, proteins which are encoded by genes positioned between *MMX123_02531* and *MMX123_02549, MMX123_02679, MMX123_03136, MMX123_03140*, and *MMX123_03143*. Many of these are identical to the maximally induced transcripts with additions of *MMX123_02679* and *MMX123_03143*. Among the proteins in the supernatant with secretion signals, 17 were detected as significantly higher abundant in xanthan and 4 as significantly higher abundant in glucose (Fig. 2F). The proteins with fold-change greater than 32-fold included four of the five genes annotated as having secreted protein products in the region *MMX123_02528* to gene *MMX123_02550* and five additional proteins MMX123_00619, MMX123_02209, MMX123_02445, MMX123_02738, and MMX123_02792.

The gene products of region *MMX123_02528* - *MMX123_02550* were consistently detected as the most induced gene products in xanthan-dependent growth with similar inductions in transcripts and proteins (Fig. S2).

### Analysis of CAZymes and putative sugar utilization loci in the genome

In accordance with the focus on xanthan utilization we inspected the genomic repertoire of *M. xanthanicum* UB-LE1 for carbohydrate active enzymes (CAZymes) and observed their response towards the different carbon sources. In total, 106 genes are annotated as CAZymes. The majority with 52 are glycoside hydrolases (GHs) followed by 39 glycosyltransferases (GTs), 5 carboesterases (CEs), 5 auxilliary enzymes, 3 polysaccharide lyases, and 1 carbohydrate binding module (CBM) protein (Fig. S3). Of these 106 annotated CAZymes, 14 CAZymes also contain a secretion signal. Seven of these belong to the GH43 family, with five significantly transcriptionally more abundant in xanthan, one significantly more abundant in supernatant protein in xanthan. One of the CAZymes with secretion signal is annotated as a putative cellulase (GH5 family) and is significantly more abundant in the extracellular proteome in xanthan. Two of the 14 CAZymes with secretion signal are increased significantly at least 32-fold on transcriptional and secreted protein level, namely a GH9 family protein and a PL8 family protein. When *pul*-loci are defined as those, which contain a GH in close proximity to a transport system consisting of periplasmic binding proteins, 15 *pul*-loci (*pul*-locus I – XV) were identified in the *M. xanthanicum* UB-LE1 genome (Table S1: pul loci). Of these, five contained at least three transcripts with significantly higher abundance in xanthan medium while not any locus contained at least three transcripts with significantly higher abundance in glucose medium. Induction in xanthan ranged from about 8-fold to up to 225-fold for a transcript in the locus with the highest induction, that is the locus *MMX123_02528 – MMX123_02550* (Table S1: pul loci).

### Identification of the *xanthan utilization* (*xut*) locus

Apparently, OMICS analyses identified a genome region between gene *MMX123_02528* to gene *MMX123_02550* as consistently upregulated during growth with xanthan as the carbon source on both, the transcript and protein level (Fig. 2, Fig. S2). A genome-wide feature analysis of GC content and transcript abundance patterns identified the same region (*MMX123_02528* - *MMX123_02550*) as having a reduced GC content compared to the surrounding regions and as carrying genes with higher transcript abundance (Fig. 3A).

**Fig. 3:**
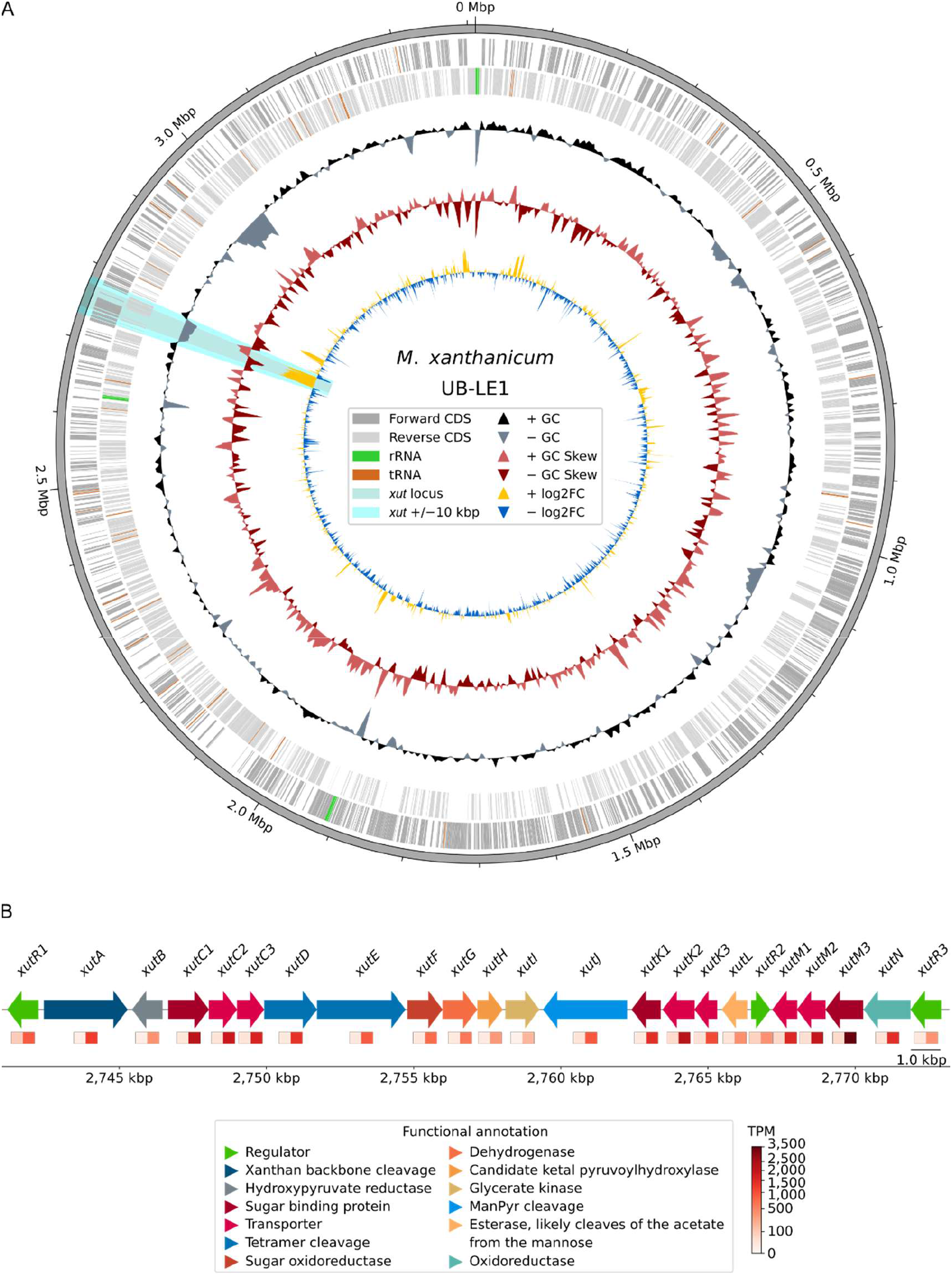
The *xanthan utilization* (*xut*) locus and its organization in the genomic context in *M. xanthanicum* UB-LE1. A) Circular genome map of the UB-LE1 genome. Track 1 displays forward coding sequences (CDS; dark gray) and track 2 shows reverse CDS (light gray). In both tracks, rRNAs are shown in green and tRNAs in brown. Track 3 represents GC content with positive GC content in black and negative in blue-gray. Track 4 depicts GC skew with positive GC skew in bright red and negative in dark red. Track 5 displays the log2 fold change (log2FC) with high log2FC in yellow and low log2FC in blue. The position of the *xut*-locus is highlighted in light teal over all tracks, with the ±10 kbp surrounding region shown in cyan. B) Organization of the *xut*-locus with its gene annotation. Genes are shown as directional arrows, representing gene orientation and their colors denote functional gene annotation. Transcript abundance for each gene is shown as two-colored rectangles below each gene. The first rectangle displays the transcripts per million (TPMs) under growth on glucose and the second rectangle represents the TPMs under xanthan growth, with increasing intensity from white to red indicating higher expression levels.

The genome analysis identified four additional regions longer than 10 kb of reduced GC content relative to the surrounding genome (Fig. 3A, Table S1: lowGC content regions) but none of these overlaps with regions of higher transcript abundance (Fig. 3A). Based on its characteristic combination of low GC content and high expression levels during growth on xanthan, genomic region 4 was designated the *xanthan utilization* (*xut*) locus (Fig. 3A, Table S1: lowGC content regions). The *xut*-locus comprises 23 genes (Fig. 3B) with functional annotations provided in Tables S1 (Table S1: xut_interproscan) and naming in Table S3. The gene *xanthan utilization regulator 1, xutR1* (*MMX123_02528*), encoded on the reverse strand, contains a DeoR-type helix-turn-helix domain and a periplasmic binding domain, suggesting a sugar-responsive transcription factor (TF). The adjacent gene *xutA* (*MMX123_02529*) encodes a secreted GH9 enzyme likely capable of cleaving the xanthan backbone following removal of terminal pyruvylated mannose residues (9). The next gene, *xutB* (*MMX123_02530*), encodes a putative 2-hydroxyacid dehydrogenase.

Genes *xutC1* to *xutI* (*MMX123_02531–02539*) form a candidate operon on the forward strand. The first three genes encode a likely ABC transporter system, including a lipid-anchored binding protein (*xutC1*) and membrane-associated transporter components (*xutC2– C3*). The intracellular enzymes XutD (GH3) and XutE (GH38) are similar to proteins, which hydrolyze glucose- and mannose-containing linkages. The downstream genes *xutF–xutH* encode proteins with predicted roles in sugar modification, including oxidoreductase and isomerase activities, while *xutI* encodes a glycerate kinase.

A second operon on the reverse strand includes *xutJ* (*MMX123_02540*), encoding a secreted PL8 xanthan lyase that likely cleaves terminal pyruvylated mannose residues, preceded by a second ABC transporter system (*xutK1–K3, MMX123_02541-02543*) and the putative esterase XutL (*MMX123_02544*). The regulator XutR2 (*MMX123_02545*), a TetR-type TF, may act as a repressor. A third operon comprises TF *xutR3* (*MMX123_02550*) and genes *xutM1–M3 (MMX123_02546-02548*), encoding a third transporter system, and XutN (*MMX123_02549*), a predicted FAD-dependent oxidoreductase. The organization of the locus suggests at least four promoters are required for full expression (Fig. 3B).

### Identification of transcriptional regulators for the *xut-*locus

The position of three TFs within the *xut*-locus suggests they may be regulating the expression of genes within and hence bind some or all the putative transcriptional starts (Fig. 3B). DNA Affinity Purification sequencing (DAP-seq) was used to determine genome-wide binding-sites for the three TF candidates XutR1, XutR2, and XutR3. The three candidates were expressed *in vitro* with halo-tags and detected on Western Blots (Fig. S4). XutR2 protein was incubated with DNA and sequencing of the bound DNA yielded two peaks with read pileups above 8,000 reads with a summit at genome position 2,766,454, which puts it 15 bases upstream of itself and 114 bases upstream of the first gene, *xutL*, in the pentacistronic transcriptional unit within the *xut*-locus (Fig. 4A).

**Fig. 4:**
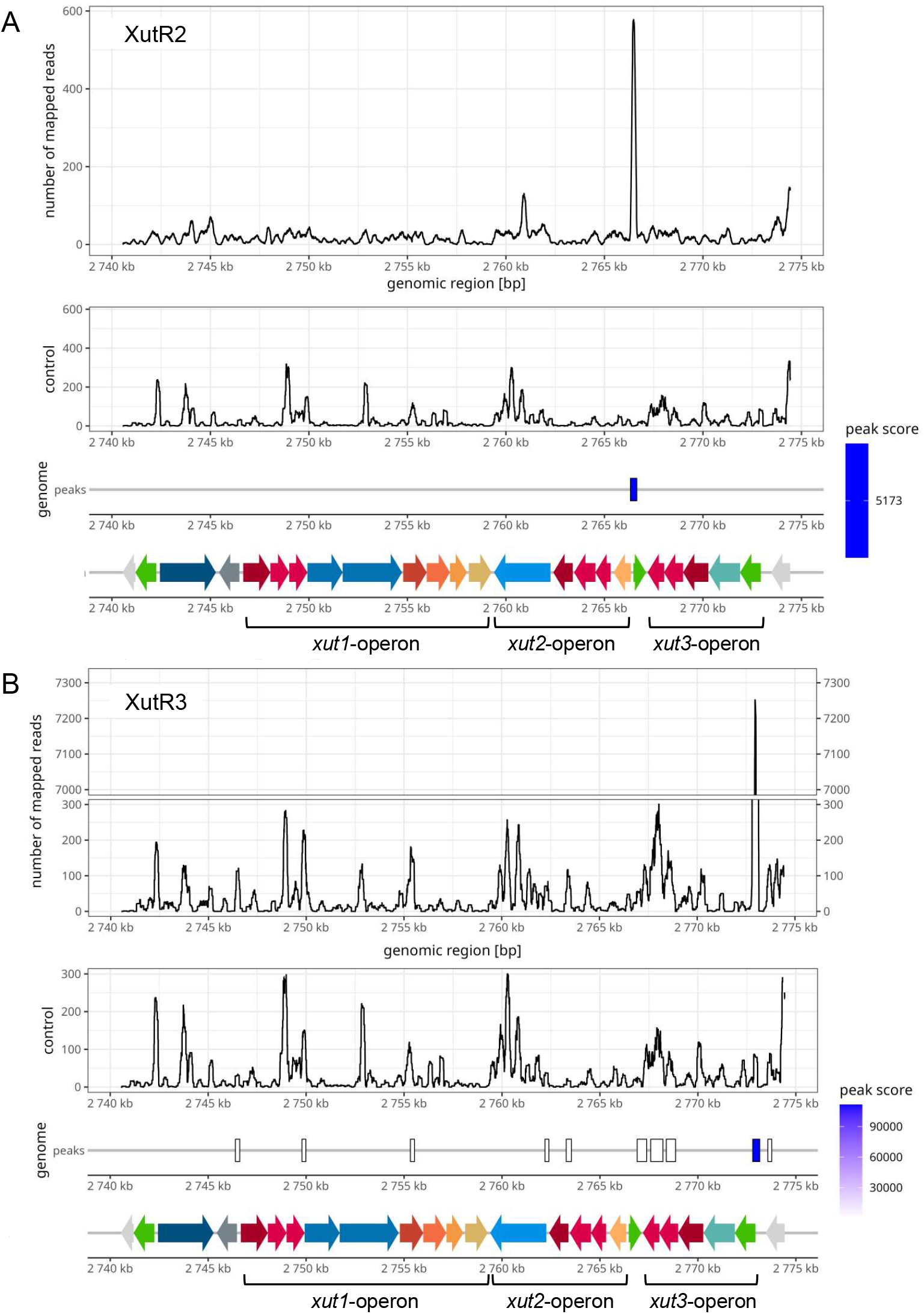
XutR2 and XutR3 bind genome regions in the *xut-*locus. DAP-seq binding data are shown for A) XutR2 and B) XutR3 across the *xut*-locus and one adjacent flanking gene on either side. For each TF, the top track depicts per-base read coverage of DAP-seq reads mapped the genome using Bowtie2. The second track shows the corresponding empty-vector control coverage, scaled to the experimental track to facilitate direct comparison of signal-to-noise ratios. The third track displays binding sites identified by MACS3 and is based on the narrowPeak output. The width of the bars indicates the called peak region, while the color intensity reflects the peak score. The bottom track illustrates schematic of the *xut*-locus genomic architecture. Arrows indicate coding strand orientation, with gene colors corresponding to the functional categories defined in Fig. 3B. Adjacent flanking genes are displayed in light gray.

The second peak is located at 1,117,528 in between *MMX123_00995* and *MMX123_00996*, neither of which is changed in transcript abundance in xanthan- or glucose-dependent growth. XutR3 yielded one peak at genomic position 2,772,971, which is 35 bases upstream of itself, the first gene in a second pentacistronic operon (Fig. 4B). Additional peaks were not detected. That is, XutR2 controls transcription of the *xut2*-operon (*xutJ* - *xutL*) and XutR3 regulates expression of the *xut3*-operon (*xutM1* – x*utR3*). Transcriptional control of the large *xut1*-operon (possibly *xutB, xutC1 - xutI*) remains elusive.

### Conservation of *xut*-loci among *Microbacterium* sp

To identify additional *Microbacterium* species carrying a *xut*-locus, proteins encoded by *MMX123_02528–MMX123_02550* were queried against 1,206 available *Microbacterium* genomes. Twelve genomes contained the core xanthan-degrading enzymes GH9 (XutA), GH3 (XutD), GH38 (XutE), and the xanthan lyase PL8 (XutJ) (Fig. 5,(10)). Three of these displayed the type I arrangement, characterized by separation of GH3/GH38 from PL8 and conservation of intervening genes, as observed in *M. xanthanicum* UB-LE1, the leaf-derived strains *M*. sp. Leaf436 and *M. testaceum* StLB037, and the wastewater-derived *M*. sp S1037. In contrast, *M. testaceum* DSM 20166 lacked the locus and was unable to grow on xanthan (Fig. S5).

**Fig. 5:**
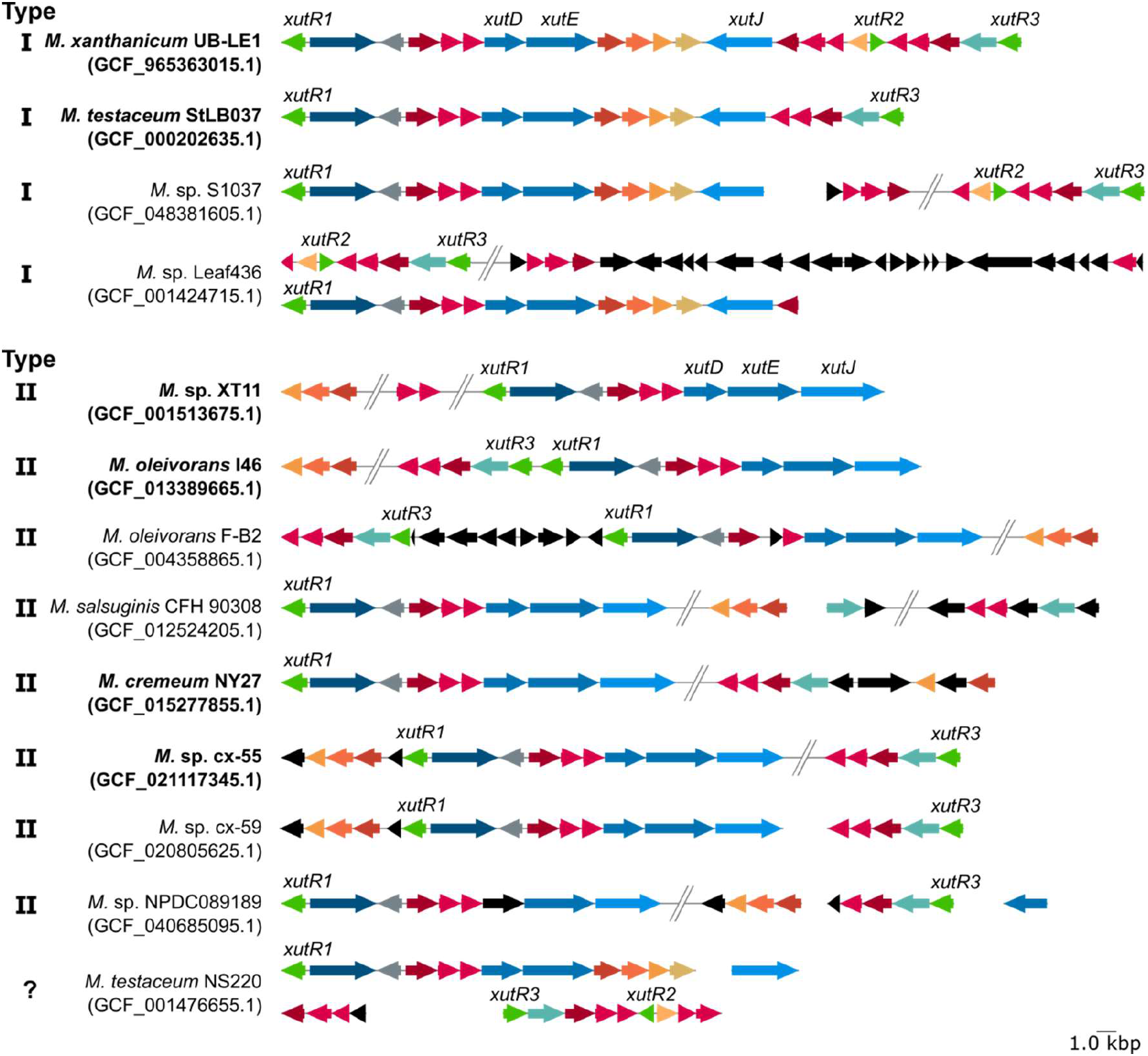
Synteny plot of genes belonging to the *xut*-locus region in different *Microbacterium* species. Color coding represents functional annotation according to *M. xanthanicum* UB-LE1 (see Fig. 3B). Chromosome-level genome assemblies are shown in bold. Genomic coordinates are listed in Table S4. Assemblies marked with Type I: Gene organization of the *xut*-locus in *Microbacterium* species with the xanthan lyase (*xutJ*) is oriented opposite to upstream genes, e.g., GH3 (*xutD*), GH38 (*xutE*). Assemblies marked with Type II: Gene organization of the *xut*-locus in *Microbacterium* species with the xanthan lyase (*xutJ*) is co-oriented with upstream genes, e.g., GH3 (*xutD*), GH38 (*xutE*). The *xut*-locus organization in *M. testaceum* NS220 could not be classified and is marked with a question mark, as the xanthan lyase (*xutJ*) is located on a separate contig from the other locus genes.

Eight genomes showed the type II arrangement, in which the genes encoding GH3, GH38, and PL8 are co-located on the same strand. Known xanthan degraders, such as *Microbacterium* sp. XT11 and fully sequenced strains including *M. oleivorans* I46 carried this organization, whereas strains lacking the locus, such as *M. oleivorans* DSM 16091 (Fig. S5), were deficient in xanthan utilization.

Across both types, the core gene block from x*utR1* to *xutE* is conserved, whereas downstream genes *xutF, xutG*, and *xutH* are variably distributed. The xanthan lyase *xutJ* is present in all strains but differs in genomic context. The second transport system and its regulator (XutR2) are restricted to type I, while the third transport system and XutR3 are consistently present in type I and variably retained in type II strains.

## DISCUSSION

The identification of a tightly regulated *xanthan utilization* (*xut*) locus in *M. xanthanicum* UB-LE1 provides insight into how soil bacteria adapt to structurally complex and spatially heterogeneous carbon sources. From an eco-evolutionary perspective, xanthan represents a particularly challenging substrate: it is chemically recalcitrant (REFs from introduction) and produced in highly localized biological contexts, for example during biofilm formation by *Xanthomonas* species (3). The strong and selective induction of the *xut*-locus in response to xanthan (Fig. 2, Fig.3), in contrast to other *polysaccharide utilization* (*pul*) loci (Table S1: pul loci), indicates that *M. xanthanicum* UB-LE1 follows a strategy of conditional investment into extracellular degradation. This strategy is consistent with the high energetic costs associated with secreting large CAZymes, such as PL8 and GH9, which release diffusible oligo- and monosaccharides(21) into the environment.

The presence of multiple, co-regulated transport systems within the same locus (Fig. 3B, Fig. 4) likely enhances substrate capture and limits loss of released sugars to competing microorganisms. Such coupling of extracellular hydrolysis with high-affinity uptake may stabilize the producer phenotype and reduce exploitation by non-degrading “cheaters” (22). The genomic potential of *M. xanthanicum* UB-LE1 further suggests a role as an early colonizer or primary degrader, analogous to pioneer degraders described in marine microbial communities that initiate polymer breakdown and structure subsequent cross-feeding interactions (23). The occurrence of *xut*-loci in leaf-associated *Microbacterium* strains (Fig. 5) indicates that xanthan degraders may also influence plant–microbe interactions by destabilizing pathogen biofilms or by redirecting carbon flow through interference competition (24). Such activity could indirectly benefit plant hosts under pathogen pressure, while simultaneously providing the degrader access to released sugars. Co-regulation of a phenylacetic acid degradation operon may further contribute to competitive fitness in biofilm-associated niches, where detoxification of antimicrobial compounds can provide a selective advantage (25). Notably, the broad repertoire of additional sugar transporters encoded by *M. xanthanicum* UB-LE1 suggests that xanthan utilization is embedded within a metabolically flexible, generalist lifestyle rather than representing a narrow specialization, consistent with the enrichment of CAZymes in plant-associated bacteria (26–28).

The genomic architecture of the type I *xut*-locus (Fig. 5) supports the view that xanthan utilization is an adaptive trait shaped by horizontal gene transfer, inducibility, and modular evolution. The reduced GC content (Fig. 3A), tight clustering of functionally linked genes (Fig. 3B), and the presence of cognate transcriptional regulators within the locus (Fig. 3B, Fig. 4) are all consistent with acquisition as a pre-assembled functional module. The organization of the *xut*-locus parallels the general logic of *pul*-loci described in Bacteroidetes (29), combining secreted and non-secreted CAZymes with tripartite transport systems. Unlike canonical Bacteroidetes *pul*-loci (30), however, the *xut*-locus includes LacI-type regulators, here designated XutR1 and XutR3 (Fig. 3B) that likely function as both sensors and transcriptional regulators via fused substrate-binding domains (Table S1: interproscan). At least one such regulator is conserved across both type I and type II *xut*-locus arrangements in *Microbacterium* (Fig. 5), suggesting functional constraints on regulatory architecture. The apparent mobility of this system is further supported by recent reports of xanthan utilization loci encoded on plasmids (17).

Comparison of type I *xut*-loci in *Microbacterium* with type II arrangements, in which xanthan-associated genes are dispersed across the genome, suggests that complete xanthan utilization requires auxiliary functions beyond the core degradative enzymes. These include enzymes, such as a 2-hydroxyacid reductase (XutB), a FAD-dependent oxidoreductase (XutN), and additional sugar-modifying activities (XutF, XutG, XutI; Table S3, Fig. 5, Fig. 6) that likely enable assimilation of unusual xanthan-derived sugars into central metabolism (Fig. 6).

**Fig. 6:**
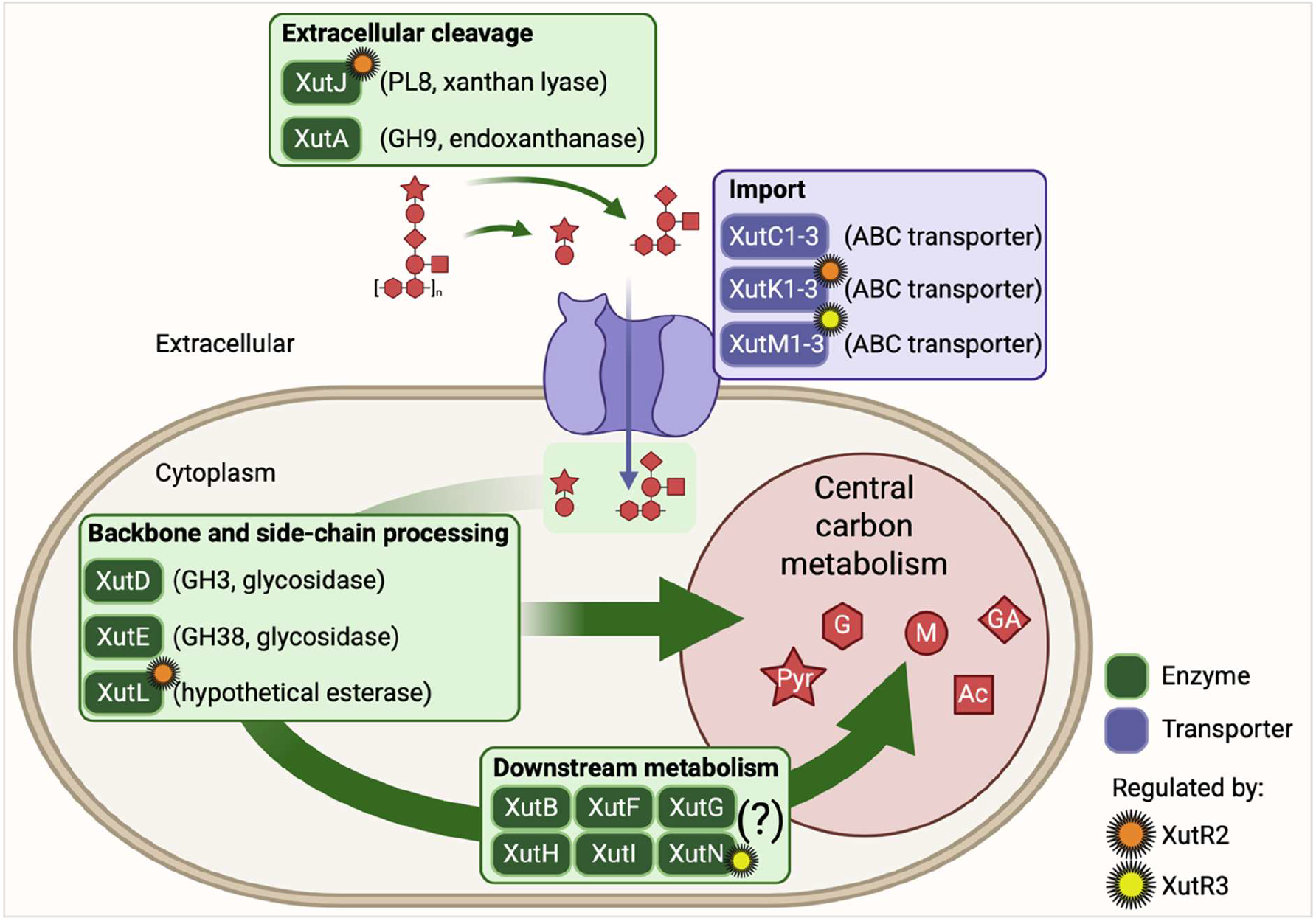
Hypothetical model of xanthan degradation and utilization (type I) in *M. xanthanicum* UB-BL1. Extracellular xanthan is initially processed by a secreted xanthan lyase (PL8 family, XutJ), which removes terminal pyruvylated mannose residues, followed by an endoxanthanase (GH9 family, XutA) that cleaves the β-(1,4)-glucan backbone into xanthan-derived oligosaccharides. These oligosaccharides as well as pyruvylated mannose are captured by surface-associated substrate-binding proteins and imported via dedicated transport systems (XutC1/2/3, XutK1/2/3, XutM1/2/3) encoded within the *xut*-locus. In the cytoplasm, an esterase (likely XutL) removes acetyl groups (Ac), and glycosidases (GH3 family, XutD and GH38 family, XutE) further hydrolyze the oligosaccharides into monosaccharides glucose (G), mannose (M), glucuronic acid (GA), which are subsequently assimilated into central carbon metabolism. The proteins XutB, XutF, XutG, XutI, and XutN are likely involved in this metabolic conversion. XutH is a candidate enzyme to cleave pyruvylated (Pyr) mannose. The question mark indicates hypothetical or unspecified functions. Expression of the xanthan utilization machinery is tightly controlled by the substrate-responsive transcriptional regulators XutR2 and XutR3 encoded within the *xut*-locus. The differently colored suns indicate transcriptional control by the respective XutR transcription factor. Protein functions are not functionally tested but suggested according to interproscan predictions.

Together, our findings position xanthan utilization as a model system for understanding how bacteria adapt to structurally complex, spatially heterogeneous extracellular carbohydrates in natural environments. The *xut*-locus in *M. xanthanicum* UB-LE1 illustrates how modular gene clusters, tight regulatory control, and metabolic integration enable conditional investment into energetically costly extracellular degradation strategies. This architecture links ecological opportunity to evolutionary innovation, highlighting how substrate availability can drive the emergence and diversification of specialized catabolic pathways. Beyond advancing fundamental insight into microbial polysaccharide degradation, our work establishes a framework for exploring and engineering xanthan-active systems. The regulatory logic and enzymatic repertoire uncovered here provide entry points for the targeted modification of xanthan structure and properties, connecting microbial ecology with future biotechnological and material applications.

## MATERIALS AND METHODS

### Isolation of *M. xanthanicum* UB-BL1 and culture conditions

Details of sample collection and strain isolation of *M. xanthanicum* UB-LE1 is described in Laker et al. (Companion paper submitted to MRA).

*M. xanthanicum* UB-LE1 was grown at 30 °C and 180 rpm in defined synthetic M9-medium (KH_2_PO_4_, 3 g L^-1^; NaCl, 0.5 g L^-1^; Na_2_HPO_4_, 6.78 g L^-1^; NH_4_Cl, 1 g L^-1^) containing 0.001 g L^- 1^ Biotin and Thiamin, 0.12 g L^-1^ MgSO_4_, 0.033 g L^-1^ CaCl_2_ and trace elements (EDTA, 0.05 g L^-1^; FeCl_3_ x 6H_2_O, 0.0083 g L-^1^; ZnCl_2_, 0.00084 g L^-1^; CuCl_2_ x 2H_2_O, 0.00013 g L^-1^; CoCl_2_ x 2H_2_O, 0.00010 g L^-1^; H_3_BO_3_, 0.00010 g L^-1^). Experiments were conducted with UB-LE1 cells, starting OD OD_600_ of 0.1, grown on 5 g L^-1^ xanthan or 6 g L^-1^ glucose to assess the influence of the carbon sources. To obtain biological replicates, three or four batches of UB-LE1 cells were grown on each carbon source.

### RNA isolation and RNA-seq analysis

Cells for RNA isolation were harvested after 34 h (+ glucose) and 48 h (+ xanthan) by centrifugation at 4 °C, 8,000 x *g* for 5 min. Pellets were frozen in liquid nitrogen and stored at −80 °C. Total RNA was extracted using the Quick RNA miniprep kit (Zymo Research, USA) following the manufacturer’s instructions. RNA quality and quantity were measured using a Bioanalyzer 2100 (Agilent, USA). rRNA was depleted using RNA Depletion llumina® Ribo-Zero Plus (Illumina, USA). Libraries were prepared using the TruSeq RNA Library Prep Kit v2 kit (Illumina, USA), pooled, and sequenced 100 bp single-end on an Illumina NextSeq 2000 instrument (Illumina, USA).

All computational analyses were performed using default parameters unless otherwise specified. Transcripts were quantified with Kallisto v0.44 (31) with parameters ‘--single -l 200 - s 20’ and the *M. xanthanicum* UB-LE1 transcript sequences as index). Downstream analyses were conducted in R v4.5.2 https://www.R-project.org/ (32) and plotted with ggplot2 v3.5.2 (33). Genes with zero counts across all samples were excluded prior to analyzes. Principle components were determined with prcomp (scale=T) and differential transcript accumulation was determined with edgeR v4.6.3 (34), classic mode) and *p*-values were corrected with Benjamini-Hochberg. A circular genome map of UB-LE1 plotted with pyCirclize (github.com/moshi4/pyCirclize) visualized gene content, GC content, and genome-wide log2-fold changes

### Protein isolation and proteome analysis

Samples were centrifuged at 8,000 x *g* for 5 min, washed with phosphate-buffered saline (PBS) containing cOmplete Protease Inhibitor (Roche, Germany), and stored at -80 °C until lyophilized. The supernatant fraction was passed through a sterile 0.2 µm filter (Fisher Scientific) and stored at -80 °C until lyophilized. Six to twelve mg of lyophilized material were resuspended in 1 mL of 100 mM ammonium bicarbonate (AmBic; Sigma-Aldrich), disrupted mechanically using a Precellys 24 homogenizer (Bertin Instruments) with 0.1 mm glass beads (BioSpec Products) at 6,500 rpm for 30 s in three cycles, refrozen in liquid nitrogen, and lyophilized.

Proteins were reduced and alkylated by sequential treatment with trifluoroethanol (VWR Chemicals), AmBic, 200 µM dithiothreitol (DTT), and 200 µM iodoacetamide (Sigma-Aldrich) for 90 min at RT, and again treated with DTT for 60 min at RT. Tryptic digestion was performed by overnight-incubation at 37 °C with sequencing-grade Trypsin Gold (1 µg µL^-1^; Promega) in AmBic and H_2_O (1:1). Peptides were purified with filter-aided sample preparation (Wisniewski et al., 2009; Sep-Pak C18 Vac Cartridges) and reconstituted in 98% LC-MS water, 2% acetonitrile, and 0.1% trifluoroacetic acid (VWR Chemicals), diluted to 0.5-1.5 µg µL^-1^. Peptides were analyzed by LC–ESI–MS/MS on a Q Exactive Plus Orbitrap instrument (Thermo Fisher Scientific), linear gradient from 3.2% to 76% acetonitrile with 0.1% formic acid over 80 min, positive ion mode from 5 to 92 min with a resolution of 70,000, AGC target of 3 x 10^6^ ions, maximum injection time (IT) of 64 ms, and a scan range of 350-2000 m/z. Data-dependent acquisition selected the top 10 precursor ions with a maximum IT of 100 ms, AGC target of 2 x 10^5^, isolation window of 1.6 m/z, and MS^2^ resolution of 17,500.

Raw data were processed using MaxQuant version 2.5.2 with default parameters except “Match between runs” enabled. Proteins were considered identified if represented by at least two unique peptides detected in a minimum of two biological replicates. For label-free quantification (LFQ), intensity values were log_2_-transformed and statistically significant differential accumulation was determined by Welch’s t-test with missing values omitted followed by Benjamini Hochberg correction (35). Secreted peptides were predicted with DeepLocPro in Gram-positive mode (36), and carbohydrate-active enzymes were annotated with dbCAN3 (37).

### Synteny analysis

The protein sequences of UB-LE1 *xut*-locus genes were queried against 1,206 downloaded proteomes from NCBI (retrieved 16.10.2025 with taxon filter “Microbacterium”; option “annotated genomes only”) using BLASTP (e-value < 1e−1, minimum sequence identity and query coverage >= 60%). Genes located 10 kbp upstream and downstream of the UB-LE1 *xut*-locus were queried with BLASTP against the proteomes with *xut*-locus proteins with *Microbacterium* sp. MM2322 as a negative control (GCF_964186585.1; (38)). The *xut*-locus regions were plotted with pyGenomeViz (https://github.com/moshi4/pyGenomeViz) with color codes based on BLASTP results.

### DNA-affinity purification sequencing (DAP-seq) and analysis

UB-LE1 was cultivated on LB agar medium over night at 30 °C. Genomic DNA was isolated using the Nucleospin Microbial DNA Kit (Macherey-Nagel, Germany, including RNase treatment), assessed on a 0.8 % agarose gel, and quantified using a High Sensitivity (HS) dsDNA assay kit on a Qubit 4 fluorometer (Thermo Fisher Scientific, USA). For DNA library preparation, 5 μg genomic DNA was fragmented, end-repaired, A-tailed, adapter ligated and amplified from 15 ng following (39).

DAP-seq experiments were conducted as described by Schiller *et al*. (2025). Briefly, coding sequences of genes *MMX123_02528* (*xutR1*), *MMX123_02545* (*xutR2*), and *MMX123_02550* (*xutR3*) were cloned in frame with an N-terminal Halo-tag encoded in the pFN19A T7 SP6 Flexi vector (Promega, USA; discontinued) using Gibson assembly (40). Primer sequences are listed in Table S5. Halo:XutR1, Halo:XutR2, Halo:XutR3, and a barnase(-) vector control were expressed with the TnT SP6 High-Yield Wheat Germ Protein Expression System (Promega, USA) using 2 µg plasmid DNA per reaction. Proteins were bound to Magnet HaloTag beads (Promega, USA) for 1 h and washed three times with a PBS-NP40 solution. Expression and bead-binding of the Halo-Tag fusion proteins were verified by western blot analysis with anti-HaloTag monoclonal antibody (Promega, USA) and Goat Anti-Mouse IgG H&L (HRP) (Abcam, UK). For DNA binding, 1 ng amplified DNA library and 1 µg salmon sperm DNA (Thermo Fisher Scientific, USA) were added to the bead-bound proteins and incubated for 1 h at 19 °C before four washes with PBS-NP40 solution and elution. The optimal number of PCR cycles for final amplification was determined by qPCR in duplicate using 1 µL recovered DNA and Luna Universal qPCR Master Mix (New England Biolabs, USA), Final libraries were amplified from 23 µL of recovered DNA using the Illumina TruSeq Strand A primer (5’-ACACTCTTTCCCTACACGACGCTCTTCCGATCT-3’) and the i7 8-baseindex primer (5’-CAAGCAGAAGACGGCATACGAGAT-[i7]-GTGACTGGAGTTCAGACGTGT-3’) with 15-19 PCR cycles. For sequencing, 5 µL of amplified DNA for each sample were pooled and purified twice with AMPure XP beads (Beckman Coulter, USA) at a bead-to-DNA volume ratio of 1:1. Library size distribution and DNA concentration were assessed using HS DNA Analysis Kit on a Bioanalyzer (Agilent Technologies, USA). and Qubit dsDNA HS Assay Kit (Thermo Fisher Scientific, USA), respectively before sequencing as single-end reads on a NextSeq2000 (Illumina, USA).

Sequencing data were processed with the automated DAP-Seq analysis pipeline v1.0.1 described in (39) using parameters ‘useDuplicates=True subsample=200000’, with the *Brassica napus* Darmor-bzh v10 genome (GCA_905183035.1,(41)) used as the random source file. Briefly, raw reads were trimmed with Trimmomatic v0.39 (42) and aligned to the *M. xanthanicum* genome (Laker et al., Companion paper submitted to MRA) using Bowtie2 v2.5.4 (43) Aligned reads were filtered by mapping quality with samtools v1.20 (44) and sambamba v1.0.1 (45) and then randomly subsampled to 200,000 per sample. Peaks were called with MACS3 v3.0.2 (46), using the corresponding empty-vector control for each TF as input control. Peaks with a fold enrichment < 3 relative to background were discarded. Genes were considered bound if a peak summit overlapped with their promoters, defined as the 150 bp region upstream of the transcription start site.

Per-base coverage across the *xut*-locus was obtained from processed Bowtie2 alignments using samtools depth and peak locations and scores were extracted from MACS3 narrowPeak output. The *xut*-locus coordinates and peak widths were displayed as gene/feature models using the R package gggenes https://wilkox.org/gggenes/ (47).

## DATA AVAILABILITY

The RNA-Seq data of the glucose and xanthan growth conditions are available in GenBank/ENA under BioProject accession number <u>PRJEB110862</u>. The GenBank/ENA accession number for the raw RNA-Seq read data under glucose are <u>ERR16928898</u>, <u>ERR16928899</u>, <u>ERR16928900</u> and under xanthan are <u>ERR16928901</u>, <u>ERR16928902</u>, <u>ERR16928903</u>.

The mass spectrometry proteomics data have been deposited to the ProteomeXchange Consortium via the PRIDE (48) partner repository with the dataset identifier PXD069319.

Strain *M. xanthanicum* UB-LE1 has been deposited in the Deutsche Sammlung von Mikroorganismen und Zellkulturen under the accession number DSM 120304 and in the Laboratorium voor Microbiologie, Universiteit Gent, collection under the accession number LMG 34097.

## ACKNOWLEDGEMENTS

We thank Ulrike Harke and Sanne Wiersma for the handling of *M. xanthanicum* UB-LE1 and the class (2023) of the master module ‘Methods and examples of functional genome research’ at Bielefeld University for experimental support. The authors would like to thank Jungbunzlauer Holding AG for their support and for providing a scholarship to fund this project, as well as for supplying “cell free” xanthan. Additionally, we would like to thank Dietmar Althaus for his aid in obtaining sampling permission. We thank the NGS team of the Bielefeld University Omics CF NGS Unit (in development) and CeBiTec as well as the technical staff of the CeBiTec Technology Platform Genomics, particularly Eva Schulte-Bernd, Yvonne Kutter, and Katharina Hanuschka for technical assistance.

## AUTHOR CONTRIBUTIONS

Bianca Laker, Computational data analysis, Manuscript writing | Michael Thomas, Strain isolation, Proteomics, Manuscript writing | Wiebke Weber, Experiments, Data analysis | Prisca Viehoever, RNA-seq, DAP-seq | Anja Meierhenrich, DAP-seq | Levin J. Klages, Genome sequencing, Computational data analysis | Tobias Busche, Genome sequencing, Computational data analysis | Karsten Niehaus, Experimental design, Supervision | Andrea Bräutigam, Experimental design, Data analysis, Supervision, Manuscript writing | Marion Eisenhut, Experimental design, Data analysis, Supervision, Manuscript writing

## ADDITIONAL FILES

The following material is available online.

## Supplemental material

**Table S1:** Additional data with mothertable, secreted CAZymes, CAZyme analysis, pul loci, low GC content regions

**Table S2:** Average nucleotide identity (ANI) values for *Microbacterium xanthanicum* UB-LE1 with closely related species.

**Table S3:** Genes of the *xut*-locus in *M. xanthanicum* UB-BL1.

**Table S4:** Genomic coordinates of sequence regions used for synteny analyses of the *xut-*locus.

**Table S5:** Primers used during initial PCR for TF candidate amplification.

**Figure S1:** Features of *M. xanthanicum* UB-LE1.

**Figure S2:** Scatterplot comparing fold-changes between transcriptome and cellular proteome.

**Figure S3:** Pie chart representing the number of proteins assigned to CAZyme classes in *M. xanthanicum* UB-LE1.

**Figure S4:** Western blotting to prove successful *in vitro* expression of XutR1, XutR2, and XutR3.

**Figure S5:** Ability of *Microbacterium* species to utilize xanthan as carbon source.

